# Antibodies targeting HSV glycoprotein B require effector functions to protect neonatal mice

**DOI:** 10.1101/2025.07.25.666806

**Authors:** Matthew D. Slein, Lesle M. Jiménez, Iara M. Backes, Evelyn M. Turnbaugh, Callaghan R. Garland, Scott W. MacDonald, Alejandro B. Balazs, David A. Leib, Margaret E. Ackerman

## Abstract

Glycoprotein B (gB) serves as the viral fusion protein for herpes simplex virus (HSV), mediating fusion between viral and host membranes resulting in infection. As such, gB represents a potentially critical target for the host immune system with high potential relevance for vaccine design. Here we investigated the mechanisms of protection for a panel of gB-specific monoclonal antibodies (mAb) in a mouse model of neonatal HSV (nHSV) infection. Viral neutralization contributed, but Fc effector functions were critical for mAb-mediated protection against nHSV mortality, depending on dose. Moreover, AAV-mediated *in vivo* expression of a gB-specific mAb in mice provided transgenerational protection against HSV-1 and HSV-2 mortality in their offspring. These findings demonstrate that antibodies targeting gB can serve as potent therapeutics and that they require diverse functional profiles to afford optimal protection, informing vaccine design.

## Introduction

Unlike many other viruses, herpesviruses split entry mechanisms across several surface glycoproteins. HSV-1 and HSV-2 have four glycoproteins that are required for viral entry^1^, glycoproteins D, B, H, and L (gD, gB, gH, gL). As these antigens are critical for infection, they are important targets for the host immune response. Viral entry begins with virus attachment to the cell surface by gB and gC, which contact heparan sulfate proteoglycans^2,3^. Virions then begin the fusion process after gD binds to one of its host cell receptors, Nectin-1 and -2, HVEM, or 3-O-sulfated heparan sulfate^1^. gD binding then triggers engagement of gB and gH/gL resulting in protein structural rearrangement between members of the fusion complex before finally leading to membrane fusion mediated by gB^4–6^. Given this activity, gB has been frequently used in past vaccine trials^7–9^ with the goal of eliciting a protective antibody response to prevent the establishment of infection.

The viral fusogen for herpesviruses, gB is a class III fusion protein^10,11^ whose ectodomain is divided into five subdomains^11^. Of these domains, DI, II, and IV are known targets of neutralizing Abs^12,13^, which presumably act by restricting rearrangement of the trimer between the metastable prefusion and lower energy postfusion states to prevent viral fusion and thus viral entry^13^. Neutralizing Abs may also function by blocking interactions between gH/gL and gB, which are required for the initiation of fusion with host cell membranes^5^. Though considered a crucial target, mechanisms of antibody-mediated protection for gB-targeting Abs remain understudied. Insights into how gB-specific mAbs mediate protection against HSV infection have the potential to drive the design of improved strategies for passive and active immunization.

Abs mediate protection through both direct and indirect antiviral activities via neutralization mediated by antigen binding and effector functions such as antibody-dependent cellular cytotoxicity, phagocytosis, and complement deposition through binding of the Fc domain to innate immune receptors. Preclinical and clinical evidence suggests that both neutralization and effector functions are important for antibody-mediated control of HSV infections^14–18^. A more complete dissection of the mechanisms by which gB-specific Abs mediate protection has yet to be performed. Specifically, defining the mechanisms of protection provides for the development of mAb therapeutics that can aid in care for vulnerable populations such as neonates and immunocompromised individuals. Although rare, neonatal HSV (nHSV) infections can have devastating consequences with high mortality rates and lifelong morbidity despite small molecule antiviral therapy^19–21^. Abs represent promising therapeutics for nHSV infections given that maternal Ab seropositivity dramatically reduces the risk of acquisition and severity of nHSV^22^.

We previously determined that mAbs targeting the viral entry mediator, gD, require neutralization and effector functions for broad and potent protection in a mouse model of nHSV infection^14^. As both gD and gB are required for viral entry, we sought to investigate whether mechanisms of Ab-mediated protection are conserved between HSV surface glycoproteins using a panel of gB-specific mAbs. By using a combination of mAbs with diverse neutralization profiles and Fc-engineering, we also sought to define the mechanisms of protection for gB-specific mAbs in a mouse model of nHSV infection. We find that gB-specific mAbs can protect neonatal mice from HSV-mediated mortality through both neutralization and effector functions, with a striking requirement for effector functions at low doses of mAb. Moreover, a gB-specific mAb expressed *in vivo* by adeno-associated virus (AAV)-vectored delivery provides transgenerational protection against neonatal disease caused by HSV-1 and HSV-2.

## Results

### Characterization of HSV gB-specific mAbs

To understand how mAbs that target gB mediate protection, we developed a panel of five distinct mAbs: Hu2c^23^, HDIT102^24^, BMPC-23^25^, Fd79^26^ and D48^27^ (**Figure 1A**). The mechanisms by which these mAbs mediate protection remain ill-defined. Moreover, these mAbs all protect adult mice from HSV-mediated mortality but have yet to be evaluated in a neonatal model of infection. Each was engineered and expressed recombinantly on a human IgG1 background. Of these, Hu2c (HDIT101) has entered clinical trials for the treatment of recurrent genital infections (NCT04165122, NCT04539483). These mAbs differ in their origin, neutralization potencies, and they target different domains of the viral fusion protein (**Figure 1A-B**). All but Fd79 have defined binding epitopes determined by structural studies^24,25,27^. Hu2c, HDIT102, and D48 bind domains critical for viral fusion (**Figure 1A-B**). Hu2c and HDIT102 both bind DI, which contains the fusion loops, and D48 binds DII, which undergoes rearrangement as a part of viral fusion^13^. On the other hand, BMPC-23 binds a region of DIV, which is presumably inaccessible in the prefusion state (**Figure 1B**), consistent with its inability to neutralize HSV.

**Figure 1:**
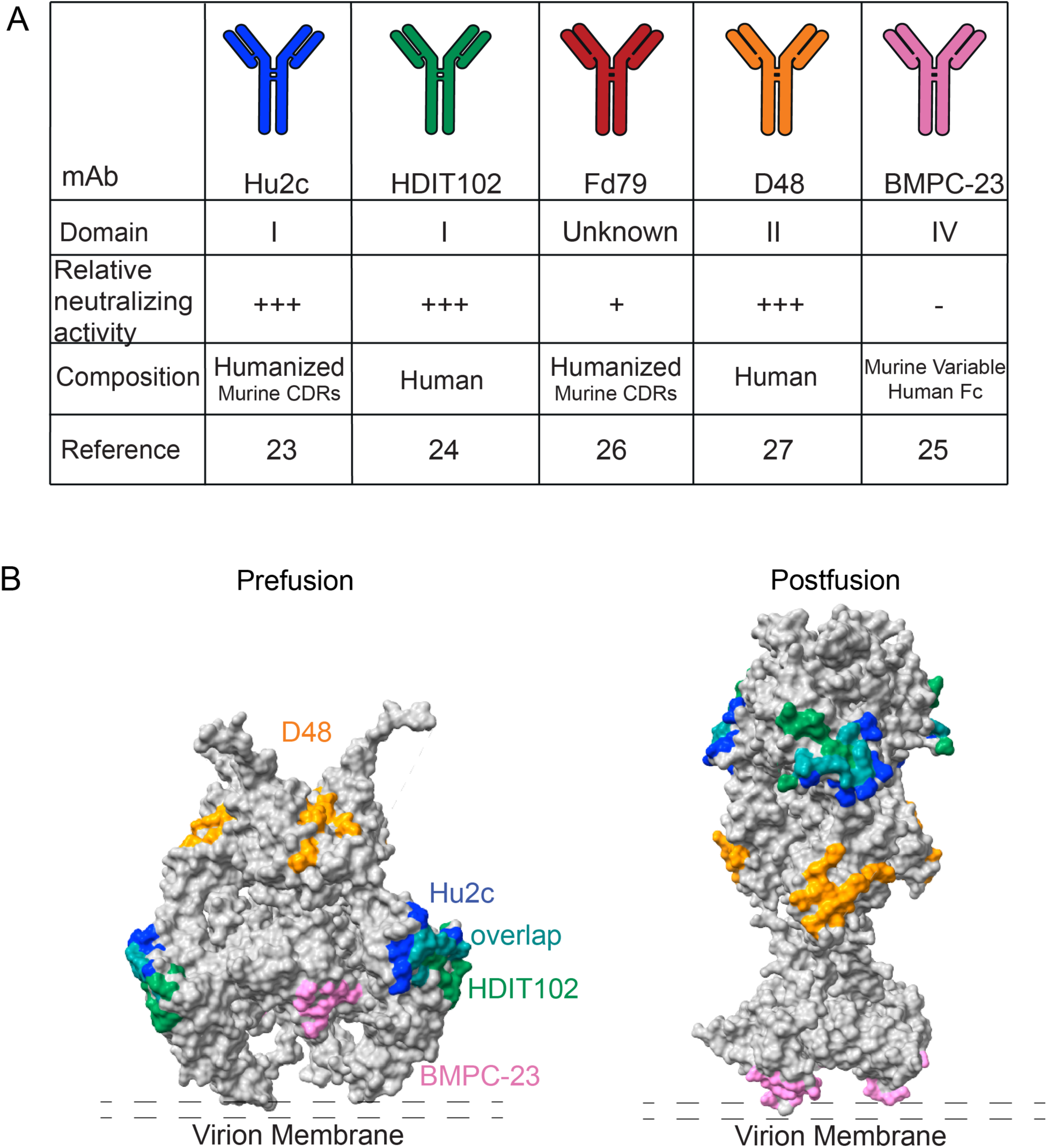
Characterization of HSV gB-specific mAbs. **A.** Table of gB-specific mAbs used in this study. Reported binding domain on gB based on available crystal structures of mAbs. Neutralization potencies for each mAb. Origin mAb composition and original reference for each gB-specific mAb **B.** Epitopes for each mAb visualized on prefusion (left, PDB: 6z9m, 41) or postfusion (right, PDB: 2gum, 42) HSV-1 gB trimers. Hu2c: blue, HDIT102: green. Overlap between Hu2c and HDIT102 – teal, D48: orange, BMPC-23: pink.

To determine mechanisms of *in vivo* protection and dissect the relative contributions of neutralization and effector functions in mediating protection against nHSV infections in a mouse model, we engineered each gB-specific mAb to lack Fc effector functions by incorporating LALA PG^28^ mutations into the Fc region. Both IgG1 and LALA PG mAbs bound to recombinant HSV-1 gB (**Figure 2A**), whereas the isotype control HSV8, an HSV gD-specific mAb, did not bind. Though the mAbs in the panel showed similar antigen recognition profiles, they differed in direct antiviral activity. Hu2c, HDIT102, and D48 all potently neutralized HSV-1 (**Figure 2B, Figure S1A**). Fd79, on the other hand, exhibited weaker neutralization potency against HSV-1 compared to the other mAbs. Regardless of mAb dose tested, BMPC-23 did not neutralize HSV (**Figure 2B, Figure S1A**). Importantly, the LALA PG versions of each gB-specific mAb retained equivalent binding and neutralization potencies compared to their respective IgG1 counterparts.

**Figure 2:**
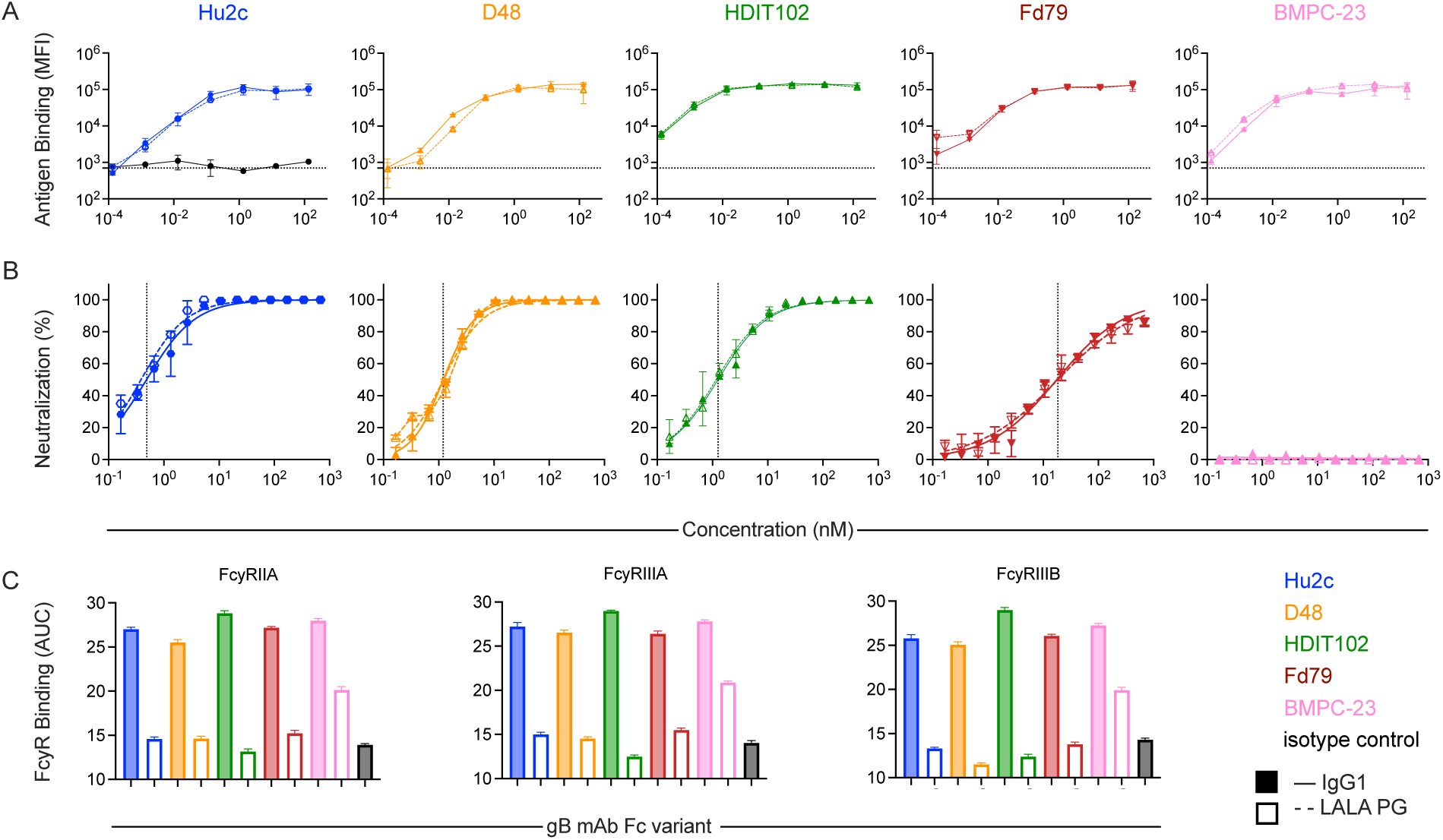
Functional characterization of WT and Fc-engineered gB-specific mAbs. **A.** Binding of each gB-specific mAb variant to recombinant HSV-1 gB via multiplex assay, and as compared to an isotype control (black, left panel). Median fluorescent intensity (MFI) is reported **B.** Neutralization activity of each gB-specific mAb against HSV-1 st17 by plaque reduction assay. Dashed vertical line indicates EC_50_ value. **C.** Human FcyR binding profiles of each gB-specific mAb variant and isotype control (black). Bar graphs represent the Area Under the Curve (AUC) values for binding to FcyRIIA (left), FcyRIIIA (center), and FcyRIIIB (right). Error bars represent standard deviation from the mean (**A-B**) or standard error of the mean (**C**). gB binding and FcyR binding experiments were performed in technical replicate. Neutralization experiments were performed in technical and 2-3 biological replicates.

Lastly, we tested the ability for the IgG1 and Fc function knockout versions of these mAbs to bind to human Fc receptors, specifically FcyRIIA, FcyRIIIA, and FcyRIIIB (**Figure 2C, Figure S1B-D**). Whereas IgG1 versions of the gB mAbs tested bound all the Fc gamma receptors tested, the LALA PG versions of all the gB mAbs had diminished binding. Intriguingly, BMPC-23 LALA PG retained some binding, suggesting context dependence to the ability of the LALA PG mutations to eliminate Fc receptor binding (**Figure S1B-D**). These results broadly indicate the suitability of the panel of mAbs to define mechanisms of *in vivo* antibody-mediated protection from nHSV when gB is targeted.

### gB-specific mAbs require effector functions to protect neonatal mice from HSV-1 infection

To examine protection from nHSV, 2-day-old C57BL/6J mice were injected intraperitoneally (i.p.) with 20 (**Figure 3A-C**), or 40 µg (**Figure 3D-E**) of the IgG1 versions of each mAb and immediately challenged with 1×10^4^ PFU of HSV-1 strain 17 (st17) intranasally (i.n). At the 20 µg dose both Hu2c and D48, which robustly neutralize HSV-1, provided nearly complete protection following HSV-1 challenge (**Figure 3A,B**). Fd79, which had a reduced neutralization potency compared to D48 and Hu2c, provided equivalent protection to the potently neutralizing mAbs (**Figure 3C**). The non-neutralizing mAb, BMPC-23, however, only protected about 40% of pups (**Figure** 3E), suggesting that effector functions contribute to mediating protection. HDIT102, despite exhibiting similar neutralization potency as D48 and binding the same epitope as Hu2c, also protected only 40% of animals (**Figure 3D**), suggesting that *in vitro* neutralization potency is not sufficient to explain the degree of protection afforded by gB-specific mAbs. In contrast, all animals treated with 40 µg of isotype control succumbed to infection (**Figure 3F**). These results indicate that while neutralization potency plays a role in mediating robust protection, it is insufficient to explain the degree of protection observed across the IgG1 mAb panel.

**Figure 3:**
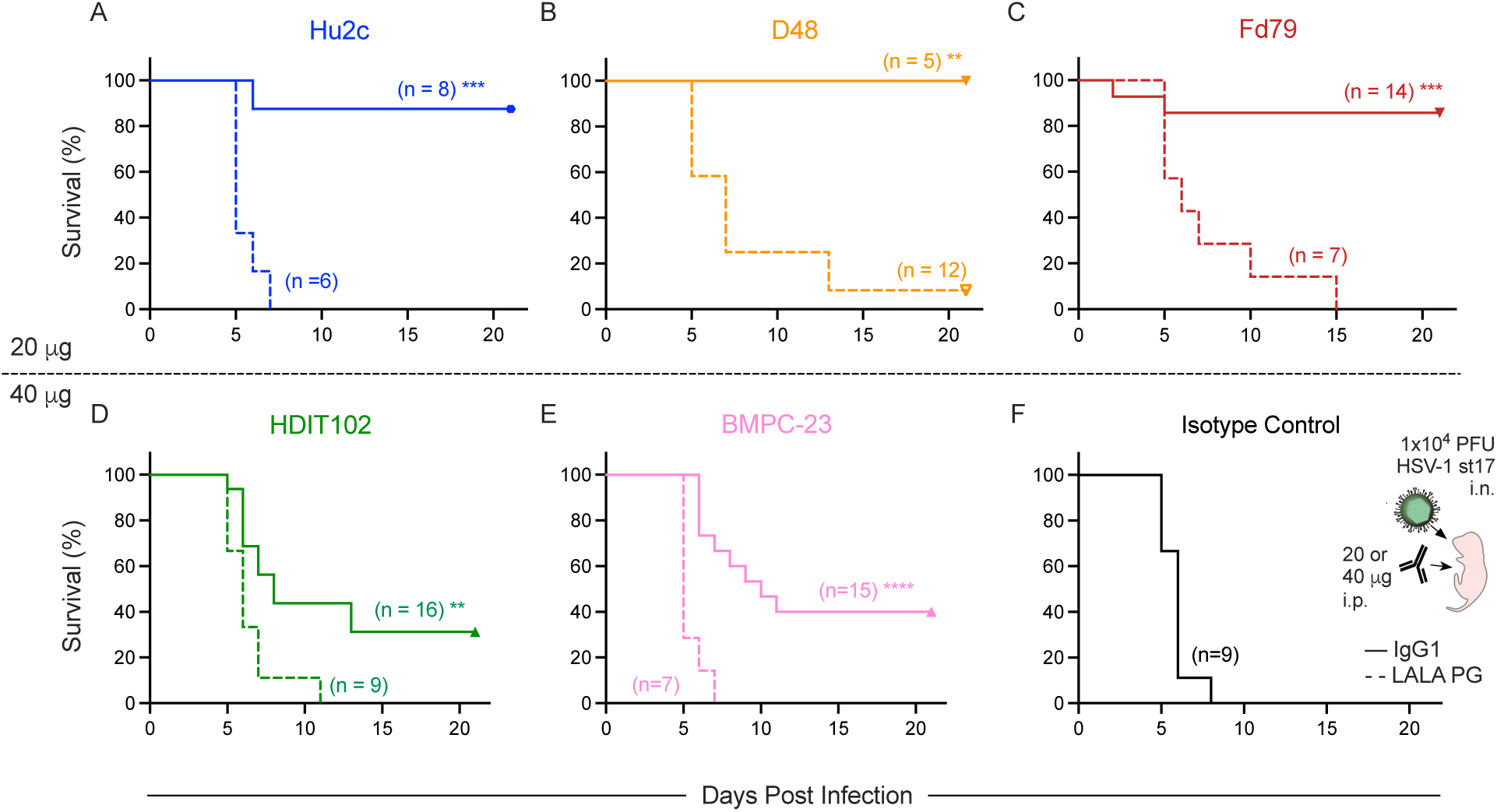
HSV gB-specific mAbs require effector functions to protect neonatal mice. **A-F.** gB-specific mAbs were delivered intraperitoneally to 2-day-old C57BL/6J mice immediately before a lethal (1×10^4^ PFU) challenge with HSV-1 st17. (**A-C**) Survival of pups given 20 µg of WT IgG1 (solid line) or LALA PG KO (dashed line) forms of Hu2c (**A**), D48 (**B**), or Fd79 (**C**). (**D-F**). Survival of pups following 40 µg of WT IgG1 or LALA PG KO mAbs. Pups were given HDIT102 (**D**), BMPC-23 (**E**) or an isotype control mAb (**F**). Number of mice in each condition are reported in inset. Statistical significance between WT IgG1 and LALA PG KO mAbs is reported in the graph as determined by the log-rank (Mantel-Cox) test. (*p < 0.05, **p < 0.01, ***p < 0.001, ****p <0.0001).

Next, to investigate whether any of these mAbs required effector functions for protection, pups were treated with the LALA PG versions of each mAb at the same doses used for their IgG1 versions. Interestingly, all mAbs, regardless of neutralization potency required effector functions to be protective in a neonatal mouse model of HSV-1 infection. These findings show that viral neutralization alone is insufficient to mediate protection against HSV-1 infection. At the same time, Fc function alone is also insufficient for complete protection, as indicated by the inability of BMPC-23 IgG1 to match the robust protection afforded by D48, Hu2c, and Fd79, even at a higher dose.

### Degree of protection depends on dose and relates to neutralization potency, but effector functions are critical even at high doses of a potently neutralizing mAb

Additional doses (10-40 µg) were evaluated for some of the mAbs (**Figure 4**). For all IgG1 mAbs tested at multiple doses, the higher dose resulted in greater survival, though testing was not always sufficiently powered to show statistical significance (**Figure 4A-C**). The degree of protection afforded by mAbs imperfectly related to their *in vitro* neutralization potency: where Hu2c exhibited the most potent activity *in vitro* (**Figure 2, Figure S1A**), D48 provided at least as much benefit *in vivo* (**Figure 4A,B**). BMPC-23, the non-neutralizing mAb, provided weak protection, even at the 40 µg dose (**Figure 4C**).

**Figure 4:**
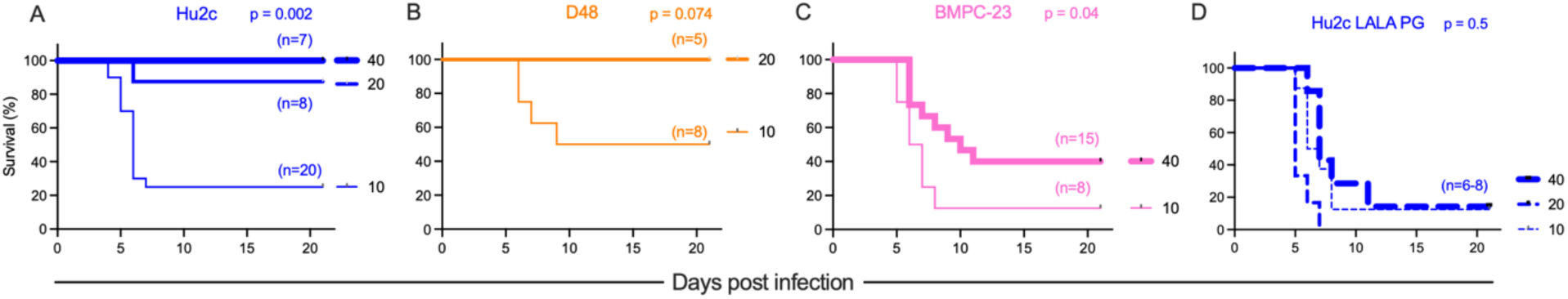
Protection relates to dose and neutralization potency but can require effector function even at high dose. gB-specific mAbs at the indicated dose (µg) were delivered i.p. to 2-day-old C57BL/6J mice prior to a lethal i.n. infection with 1×10^4^ PFU of HSV-1. Survival of mice following treatment with Hu2c IgG1 (**A**), D48 IgG1 (**B**), BMPC-23 IgG1 (**C**), or Hu2c LALA PG (**D**). Number of mice in each condition is reported in inset. Statistical significance between between highest and lowest doses tested as determined by the log-rank (Mantel-Cox) test.

We previously demonstrated that mAbs specific for gD can exhibit dose-, route-, and virus strain-dependent impacts on mechanisms of protection^15,29^. To begin to explore whether a high mAb dose could overcome the requirement for mAb effector function observed for this panel of gB-specific mAbs, Hu2c was tested in LALA PG form across the full dose series. In contrast to its IgG1 form, Hu2c LALA PG did not demonstrate dose-dependent protection (**Figure 4D**), suggesting that increasing dose over this range was insufficient to compensate for the loss of binding to FcR, resulting in an inability of this potently neutralizing mAb to protect from nHSV.

### Robust cross-strain protection afforded by AAV-expression of even a poorly neutralizing mAb

Having established that direct administration of gB-specific mAbs to pups resulted in dose-dependent protection, we wanted to address whether alternative mAb delivery systems could also be efficacious and if protection across serotypes could be observed. The neutralization efficiency of these gB-specific mAbs was evaluated for both HSV-1 st17 and HSV-2 strain G (stG) (**Figure 5A-B**). Genes to express Fd79 in IgG1 form were administered by vectored immunoprophylaxis (VIP), in which an HSV-specific mAb is expressed and secreted from muscle cells *in vivo* following adeno-associated virus (AAV)-vectored delivery. Female mice received one intramuscular injection of a nonintegrating AAV8 vector encoding Fd79 (**Figure 5C**). *In vivo* expression was confirmed by collecting sera weekly from transduced mice (**Figure 5D**). All four dams had robust and stable expression of Fd79 *in vivo* for at least one month (**Figure S4**). We then assessed whether expressed mAb was trans-generationally transferred and protective to offspring against HSV challenge, mimicking the protection afforded by maternal HSV seropositivity in humans against neonatal disease. Offspring of AAV-treated dams were challenged with 1×10^4^ PFU of HSV-1 st17 or 300 PFU of HSV-2 stG on day 2 of life. All the progeny of Fd79 AAV-transduced dams were completely protected from HSV-1- and HSV-2-mediated mortality (**Figure 5E**). These findings indicate that *in vivo* expression of Fd79 and subsequent transfer to pups is robustly protective and therefore represents an alternative option to single bolus mAb delivery to protect at risk neonates from HSV infection. Together these results indicate that diverse gB-specific mAbs protect against nHSV infection and that degree of protection depends on mAb dose, viral neutralization potency, and antibody effector functions.

**Figure 5:**
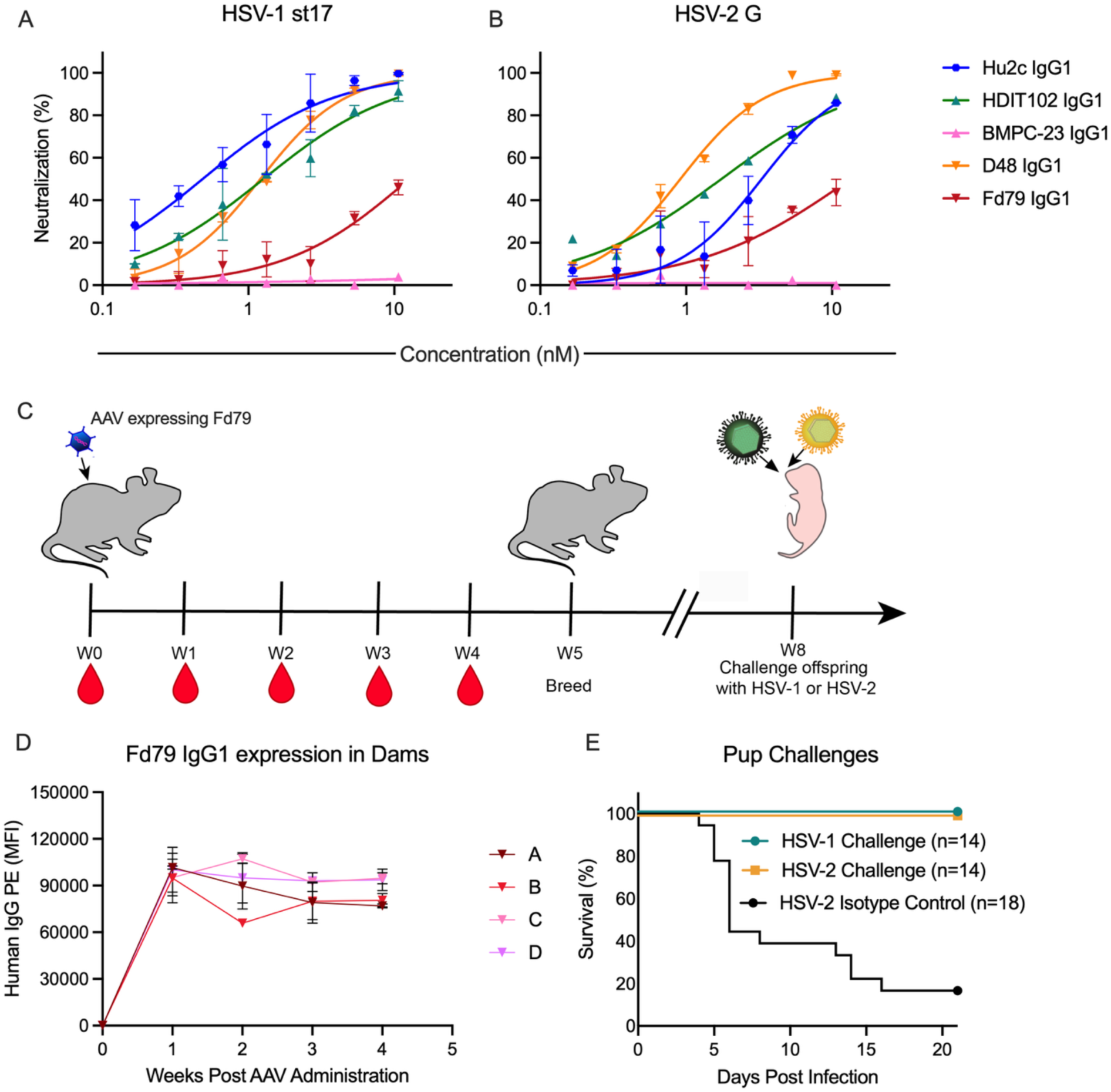
Broad intergenerational protection can be accomplished through vectored delivery of a gB-specific mAb. **A-B.** Neutralization potencies of gB-specific mAbs against HSV-1 st17 (**A**) and HSV-2 G (**B**). **C.** Experimental overview and study design in which AAV encoding the gB-specific mAb, Fd79, on a human IgG1 backbone was administered intramuscularly to 4 female mice prior to breeding and viral challenge of pups. **D.** Expression of AAV-expressed Fd79 in the sera of female mice at weeks 0 through 4 post AAV administration. **E.** Survival of offspring of AAV-treated dams challenged with 1×10^4^ PFU of HSV-1 st17 or 300 PFU of HSV-2 G on day 2 postpartum.

## Discussion

Antibodies as therapeutics represent promising drugs for the prevention and treatment of viral infections, especially when efficacious vaccines are unavailable. Determining the dominant mechanisms of Ab-mediated protection is a critical step in the design and optimization of potential mAb therapies. In this study, mAb mechanism was dependent on dose, effector functions, and viral neutralization. While viral neutralization likely contributes to mAb-mediated protection, the ability for gB specific mAbs to mediate Fc effector functions was unexpectedly crucial. Hu2c, which potently neutralizes HSV-1, required Fc effector functions to be efficacious at all doses tested. Fd79 poorly neutralizes HSV-1 *in vitro* but provided comparable protection to Hu2c *in vivo* and similarly required effector functions to be protective. HDIT102, which neutralizes HSV-1 with potency intermediate to Hu2C and Fd79, failed to protect most pups following challenge with HSV-1. D48, like Hu2c, also required effector functions to protect neonatal mice from HSV-mediated mortality. These results indicate that neutralization potency may not be the dominant mechanism of protection even when the mAb is present at high doses. In contrast, BMPC-23, the non-neutralizing mAb, only mediated partial protection even at the highest dose tested – indicating that Fc function alone is only partially protective for this gB-specific mAb.

As we have observed previously^14,15,29^, Ab dose was a critical determinant for understanding the dominant mechanism(s) of protection against HSV-mediated mortality. We saw an apparent dose-dependent increase in protection for protection against nHSV mortality for the direct administration of mAb to pups. At the lowest dose tested (10 µg/pup), none of the IgG1 mAbs tested were able to fully protect pups from HSV-1-mediated mortality, but protection was improved by increasing the mAb dose. Despite the dose-dependent increase in protection afforded by Hu2c IgG1, the consistent inability for Hu2c LALA PG to provide protection indicated that Hu2c requires effector functions to be protective even when given at a high dose. These findings differ from our previous work with gD-specific mAbs^15^, where viral neutralization was the dominant mechanism of protection at high mAb doses and effector functions were only required at “subneutralizing” mAb doses. It is likely that different glycoprotein targets may be variably susceptible to different mechanisms of Ab-mediated protection, which is critical for both therapeutic and vaccine design. Together these results indicate that direct administration of gB-specific mAbs represents a potent approach to protect vulnerable neonates.

While showing their importance, our data do not enable us to pinpoint the relative contributions of different effector functions. The LALA PG mutant abrogates binding to all FcgR and to C1q, imparting a global reduction in effector function. Other experiments, such as with targeted knockouts of different cell types, the complement cascade, or individual receptors, would be needed to parse the contributions of each distinct effector function. Such experiments were not pursued, as evidence from mouse models^30^ and humans^31^ suggests that the balance of different functions will differ in association with host genetics. Our work with gD-specific mAbs also shows that this balance can differ between virus strains. In this context, further refinement of mechanism is expected to impact the generalizability of refined insights.

Other limitations include only testing mAb efficacy against a single laboratory strain of HSV-1, which may not fully recapitulate virulence and pathogenesis of circulating HSV strains^32^. Moreover, we focused on HSV-1 in the direct administration experiments as it has become a major cause of neonatal disease in the US^33^ and has a global seroprevalence of 66%^34^ indicating a high burden of disease. Whether our findings translate to HSV-2 will have to be determined. While mice are a useful model system to perform preclinical experiments, they differ from humans in numerous ways, especially in regards to FcyR distribution and expression on immune cells^35^. These differences may impact clinical translation and the importance of different Ab-mediated mechanisms of protection in humans.

Beyond direct administration of mAbs, we also demonstrated prolonged delivery of mAb to pups following VIP treatment of dams. Here, we observed that Fd79 is efficiently expressed *in vivo* and that its maternal transfer protects offspring from mortality caused by HSV-1 and -2. In this system, mAb is continuously expressed and transferred across the placenta prior to birth to distribute throughout the pup^14^, mimicking the protection afforded by maternal seropositivity in humans^22^ and maternal vaccination strategies in mice^36^. Moreover, unlike in the direct administration experiments in which mAbs are only delivered once, pups are likely continually supplied with AAV-expressed mAb through nursing, which is a robust mechanism of Ab transfer in rodents^37^.

While we observed a clear role for Fc function in mediating protection against nHSV infection, the relatively minor role for viral neutralization was surprising. This data potentially aligns with failed vaccine trials for HSV in humans. A major vaccine trial that used gD-2 and gB-2 as targets induced robust neutralizing responses but Abs with little to no Fc functional capacity^38^ ultimately failed to meet efficacy criteria and advance beyond clinical trials. Our findings in understanding Ab mechanism of action against various HSV antigen targets has demonstrated the importance of Fc effector functions for protection^15,29^. It is possible that vaccine strategies that robustly induce effector function responses like ADCC, in addition to viral neutralization, will be more efficacious against HSV infections. Existing preclinical and clinical data suggest the importance of vaccine strategies for HSV that induce effector functions^7,39–41^.

In summary, this study demonstrated that administration of HSV gB-specific mAbs, through either recombinant protein or vectored delivery, potently protect neonatal mice from HSV-mediated mortality. Efficacy was directly influenced by Ab dose, neutralization potency, and Fc effector functions, demonstrating that mAbs with polyfunctional profiles are promising candidates for HSV treatment and prevention, particularly for neonatal infections.

## Materials and Methods

### Mouse Procedures and Viral Challenges

C57BL/6J (B6) mice were either purchased from The Jackson Laboratory or bred in house in accordance with Institutional Animal Care and Use Committee approved protocols. mAbs were administered via the intraperitoneal route to 2-day-old pups via a 25 µL Hamilton syringe in a 20 µL volume under 1% isoflurane anesthesia. The viral strains used in this study were HSV-1 st17syn+^42^ and HSV-2 G^43^ (provided by Dr. David Knipe). Viral stocks were prepared using Vero cells as described elsewhere^44,45^. Newborn pups were infected intranasally (i.n.) on day 2 of life with 1×10^4^ PFU of HSV-1 st17 or 300 PFU of HSV-2 stG in a 5 µL volume under 1% isoflurane anesthesia. Pups were then monitored and weighed daily through day 21 post infection. Endpoint criteria for the viral challenge experiments were defined as excessive morbidity (hunching, spasms, and/or paralysis) and/or >10% weight loss from the previous measurement. All animal procedures were performed in accordance with Dartmouth’s Center for Comparative Medicine and Research policies and following approval by Dartmouth’s Institutional Animal Care and Use Committee (Protocol number: 00002151 – approved 240809).

### Monoclonal Antibodies

Hu2c^23,24^, HDIT102^24^, Fd79^26^, BMPC-23^25^, and D48^27^ VH and VL sequences were collected from published sequences or published structures and synthesized by Twist Biosciences as full length IgG1 heavy chains and Kappa (Hu2c, Fd79, BMPC-23 and D48) or Lambda (HDIT102) constant light chains. Recombinant DNA was cloned into pCMV expression vectors. Fc variants of each gB-specific mAb were generated via overlap extension PCR combining the VH for each mAb with constant domains containing the specific Fc mutations. All expression plasmids containing mAb heavy or light chain sequences were sequence confirmed via Sanger or whole plasmid sequencing (Azenta/GENEWIZ). mAbs were expressed via the co-transfection of heavy and light chain sequences in ExpiCHO cells (ThermoFisher) in accordance with manufacturer’s protocols. At 8-10 days post transfection, cultures were spun for 3 hours at 3000 RCF to pellet the cells. Supernatants were then sterile filtered (0.22 µm). All mAbs were purified using custom packed protein A columns (Cytiva) or protein A/G plus agarose (ThermoFisher) and eluted with 100mM glycine pH 3.0. Eluates were immediately neutralized using 1M Tris-HCl pH 8.0. mAbs were concentrated and buffer exchanged into PBS using Amicon 30kDa cut off filters. mAbs were passed over endotoxin removal columns, aliquoted, and snap-frozen before being stored at −80°C until use. ChimeraX^46^ was used to make representative figures with the known epitopes of the gB-specific mAbs used in this study using the determined structures of HSV-1 pre^47^ and postfusion^48^ gB.

### Measurement of binding to HSV-1 gB

Recombinant HSV-1 gB antigen (provided by Dr. Gary Cohen) was coupled to MagPlex (Luminex) beads as previously described^49^. gB-specific mAbs were serially diluted in 1x PBS with 0.1% bovine serum albumin (BSA) and 0.05% Tween-20 and incubated with antigen-coupled beads overnight at 4°C with constant shaking. Antibody immune complexes were washed once before being incubated with an anti-human IgG-PE secondary antibody (Southern Biotech) for 1 hour at room temperature with constant shaking. Beads were washed a second time and then analyzed on the xMap system (Luminex). The median fluorescence intensity of at least 10 beads/region was recorded. An isotype control antibody and buffer-only well control were used to determine assay background.

### Measurement of antibody binding to human Fc receptors

Serially diluted gB-mAbs were incubated with gB and anti-human IgG MagPlex beads as described above. Beads were washed before being incubated with recombinant biotinylated human Fc gamma receptors^50^ (Duke Human Vaccine Institute) that were tetramerized with streptavidin-PE for 1 h. The beads were washed and analyzed on the xMap system. The median fluorescence intensity of at least 10 beads/region was recorded. An isotype control antibody and a buffer only control were used to determine antigen-specific binding and assay background signal. Area under the curve was calculated using Prism 10 (GraphPad).

### Viral Neutralization

Viral neutralization of HSV-1 and HSV-2 was performed via plaque reduction neutralization as previously described^15,29^. Briefly, serially diluted mAbs were incubated with 100 PFU of HSV-1 st17 or HSV-2 stG for 1 hour at 37°C before being added to confluent vero cell monolayers grown in 6 well plates (Corning). Virus:mAb immune complexes were incubated with cells for 1 hour at 37°C with shaking every 15 minutes. After the incubation, methylcellulose overlay was added to the cells. Plates were incubated for 48 (HSV-1) or 72 (HSV-2) hours at 37°C with 5% CO_2_. After incubation, the overlay was removed, and cells were fixed with 1:1 methanol:ethanol for 30 minutes at RT. The plates were then stained with 12% Giemsa overnight. The stain was removed, and plaques were counted on a light box. Viral neutralization (%) was calculated as [(number of plaques in virus only control well - number of plaques counted at mAb dilution)/# of plaques in virus only control well]*100. Assays were performed in technical and 2-3 biological replicates.

### AAV Production and Mouse Procedures

AAVs encoding the heavy and light chain sequences of Fd79 on a human IgG1 backbone were constructed as previously described^51^. A single 40 µL injection of 1×10^11^ genome copies of the AAV was administered into the gastrocnemius muscle of female B6 mice as previously described^14^. Dams were bled weekly via the mandibular vein with a 5mm lancet. Blood was allowed to clot by stasis at room temperature for 30 minutes and then spun at 2000 RCF for 20 minutes at 4°C. Sera was collected and stored at −20°C. AAV-driven antibody expression was verified using the magnetic bead based assay as described above. Serially diluted sera were incubated with gB and anti-human IgG capture beads and detected with an anti-human IgG-PE secondary antibody (Southern Biotech) and analyzed via the xMap system (Luminex).

## Acknowledgements

We thank all members of the Ackerman and Leib labs for their scientific insights and experimental guidance. Molecular graphics and analyses performed with UCSF ChimeraX, developed by the Resource for Biocomputing, Visualization, and Informatics at the University of California, San Francisco, with support from National Institutes of Health R01-GM129325 and the Office of Cyber Infrastructure and Computational Biology, National Institute of Allergy and Infectious Diseases.

## Author Contributions

M.D.S., D.A.L., and M.E.A conceptualized the study. M.D.S., L.M.J., I.M.B., E.M.T., C.R.G., performed experiments. S.W.M. and A.B.B produced and provided AAV. M.E.A. and D.A.L. obtained funding and supervised the work. M.D.S. drafted figures and wrote the manuscript. M.D.S., D.A.L., and M.E.A. finalized the manuscript. All authors have read and edited the final version of the manuscript.

