## Supplemental Materials for "Antibodies targeting HSV glycoprotein B require effector functions to protect neonatal mice"

### Supplemental Figures and Legends

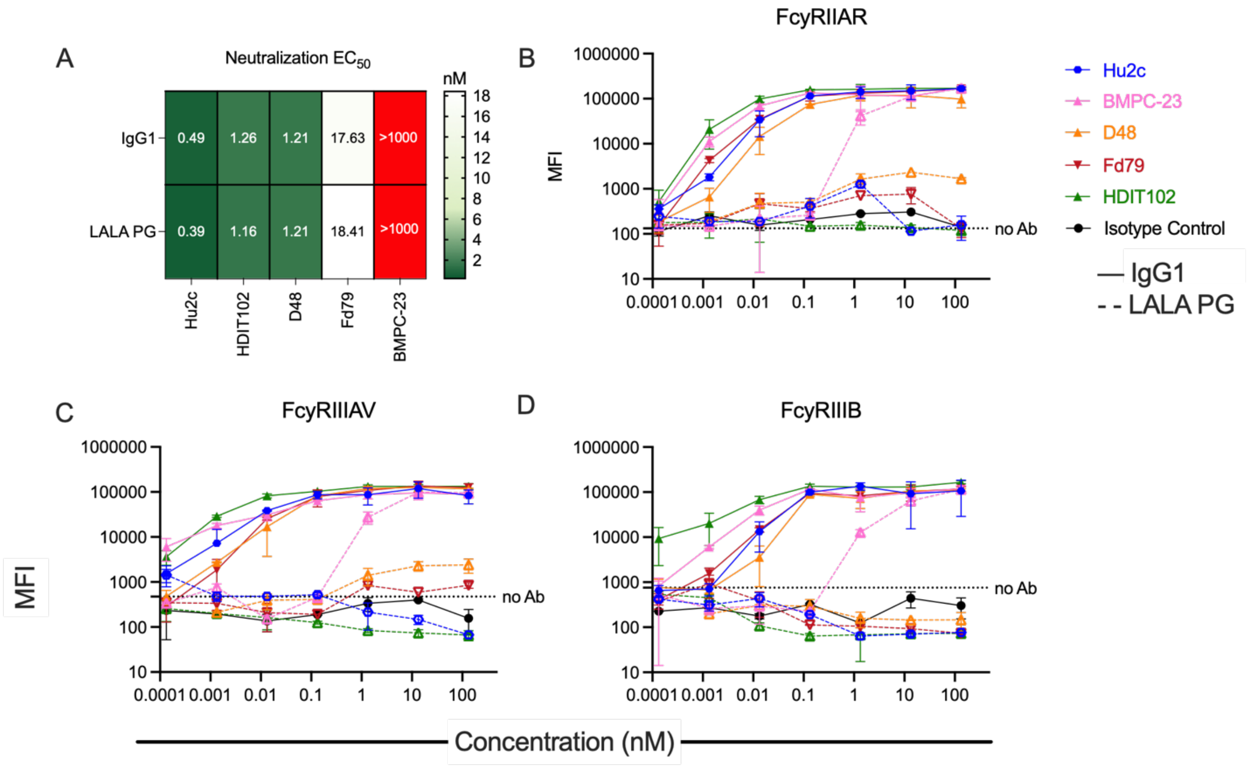


**Supplemental Figure 1 (Related to Figure 1): Neutralization potencies and FcγR binding of gB-specific mAbs and their Fc-engineered variants. A**. Heatmap of neutralization midpoint Effective Concentration ($EC_{50}$) values for HSV-1 between the IgG1 and LALA PG Fc variants of the gB-specific mAbs. **B-D**. Median fluorescent intensity (MFI) of tetramerized human FcγRIIA (**B**), FcγRIIIA (**C**), or FcγRIIIB (**D**) binding to each indicated HSV-specific mAb when bound to gB.

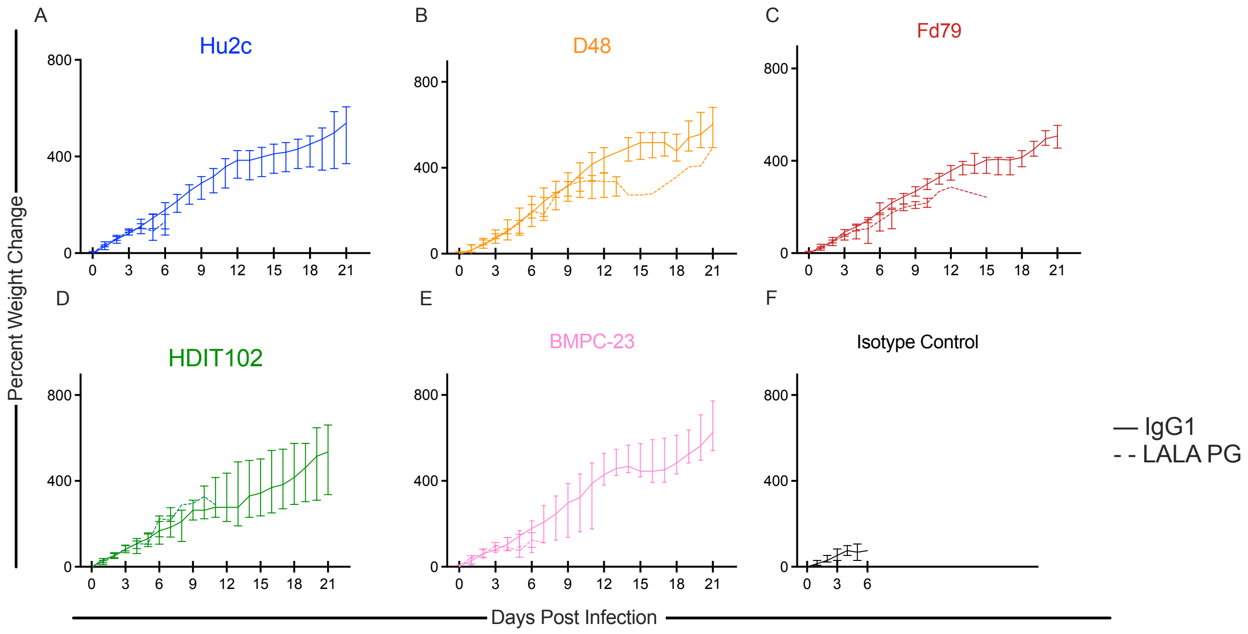


**Supplemental Figure 2 (Related to Figure 3) Percent weight gain of mice following HSV-1 challenge.** Mice were weighed daily for 21 days post challenge and treatment with indicated mAb at either 20 (top row) or 40 (bottom row) µg dose. Error bars represent the standard error of the mean.

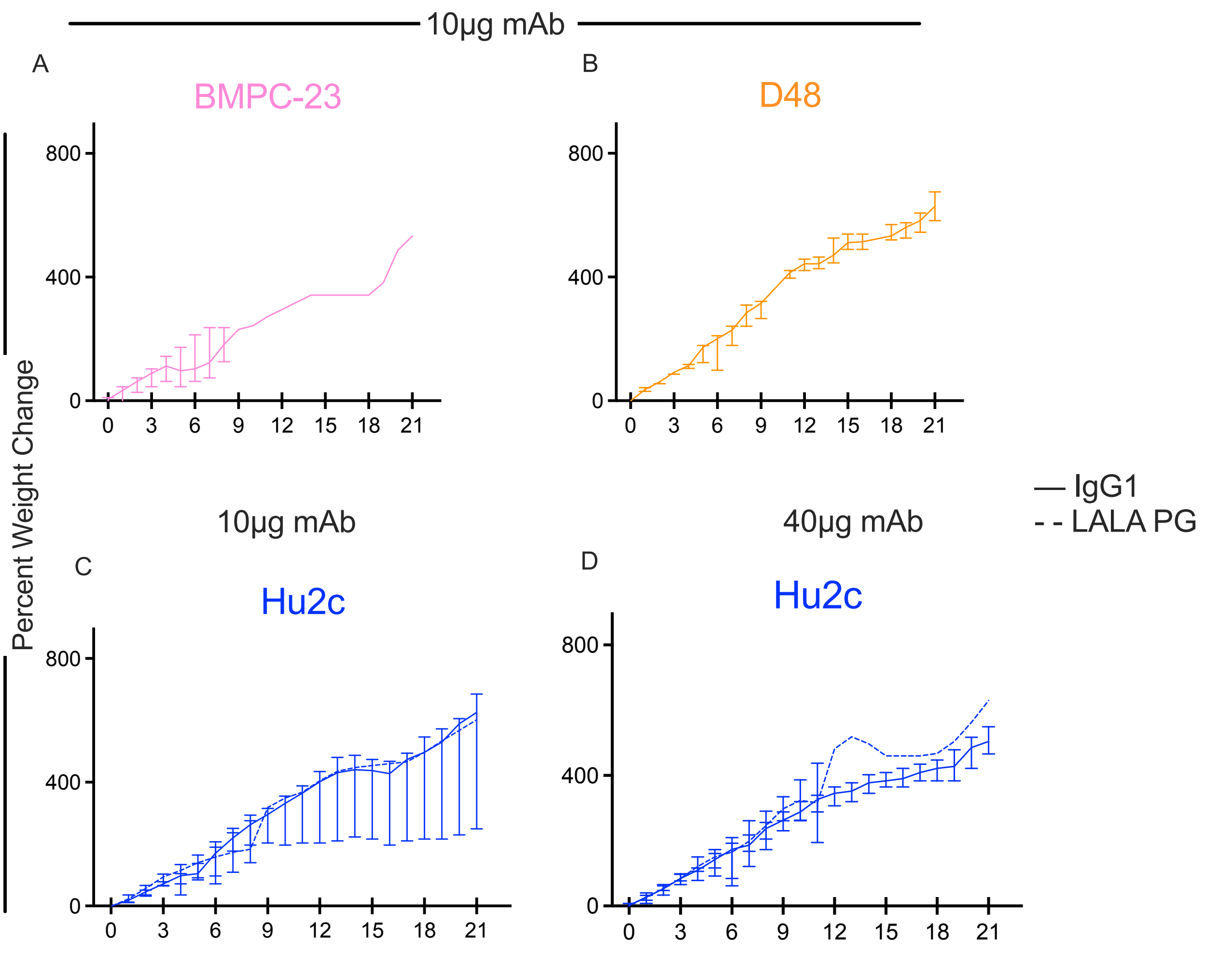


**Supplemental Figure 3 (related to Figure 4)**. **Percent weight gain of mice given alternative doses of gB-specific mAbs following HSV-1 challenge.** 2-day-old C57BL/6J mice received 10 (**A-C**) or 40µg (**D**) of the indicated gB-specific mAbs or isotype control delivered i.p. immediately before a lethal challenge with $1x10^4$ PFU of HSV-1 st17. Mice were monitored for 21 days post infection and weighed daily. Error bars represent the standard error of the mean.

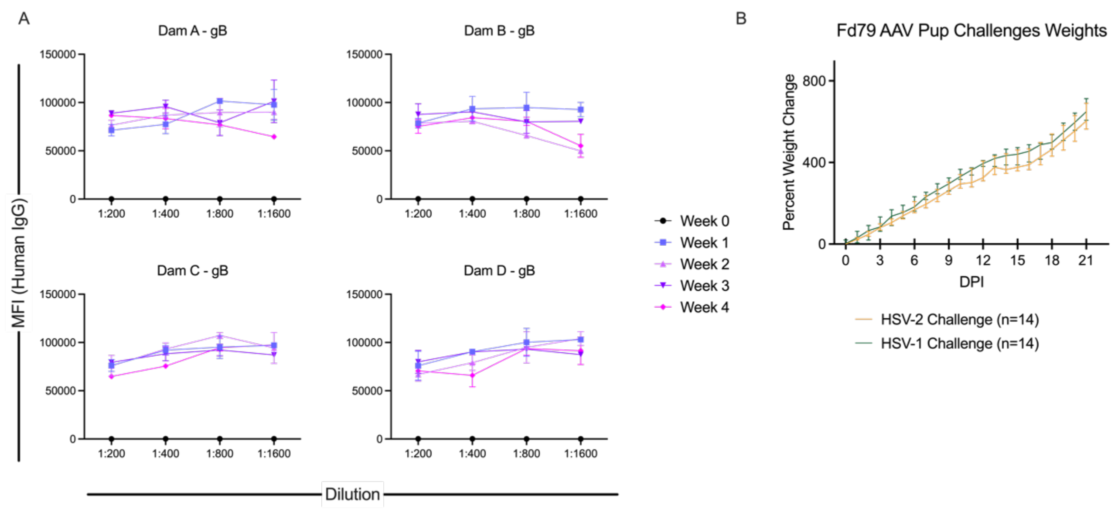


**Supplemental Figure 4 (related to Figure 5) AAV-expression of Fd79 is stable over time and protects offspring from HSV-mediated morbidity and mortality. A**. *In vivo*-expressed Fd79 was detected in the sera of 4 female mice at weeks 0-4 post transduction. Error bars represent standard deviation from the mean **B**. Percent weight gain of offspring of Fd79-AAV-transduced dams following HSV-1 or HSV-2 challenge. Error bars represent standard error of the mean.
